# Caspase-mediated cleavage of murine norovirus NS1/2 potentiates apoptosis and is required for persistent infection of intestinal epithelial cells

**DOI:** 10.1101/372615

**Authors:** Bridget A. Robinson, Jacob A. Van Winkle, Broc T McCune, A. Mack Peters, Timothy J. Nice

## Abstract

Human norovirus (HNoV) is the leading cause of acute gastroenteritis and is spread by fecal shedding that can often persist for weeks to months after the resolution of symptoms. Elimination of persistent viral reservoirs has the potential to prevent outbreaks. Similar to HNoV, murine norovirus (MNV) is spread by persistent shedding in the feces and provides a tractable model to study molecular mechanisms of enteric persistence. Previous studies have identified non-structural protein 1 (NS1) from the persistent MNV strain CR6 as critical for persistent infection in intestinal epithelial cells (IECs), but its mechanism of action remains unclear. We now find that the function of CR6 NS1 is regulated by apoptotic caspase cleavage. Following induction of apoptosis in infected cells, caspases cleave the precursor NS1/2 protein, and this cleavage is prevented by mutation of caspase target motifs. These mutations profoundly compromise CR6 infection of IECs and persistence in the intestine. Conversely, NS1/2 cleavage is not required for acute replication in extra-intestinal tissues or in cultured myeloid cells, indicating an IEC-specific role. Intriguingly, we find that caspase cleavage of NS1/2 reciprocally promotes caspase activity, potentiates cell death, and amplifies spread among cultured IEC monolayers. Together, these data indicate that the function of CR6 NS1 is regulated by apoptotic caspases, and suggest that apoptotic cell death enables epithelial spread and persistent shedding.

**Author Summary:** Human Norovirus infection is highly contagious and the most common cause of acute gastroenteritis. Norovirus can be persistently shed after resolution of symptoms, perpetuating or initiating new outbreaks. Murine norovirus (MNV) is also persistently shed, enabling study of host and viral determinants of norovirus pathogenesis. We previously identified a critical role for MNV non-structural protein 1 (NS1), in persistence. Herein we find that regulation of NS1 by host apoptotic caspases is required for infection of intestinal epithelial cells, but not for extra-intestinal spread. Additionally, we demonstrate that NS1 reciprocally promotes cell death and spread among epithelial cells. These data identify regulation of NS1 by host proteases and suggest that apoptotic death is a determinant of epithelial spread and persistence.

## Introduction

Human norovirus (HNoV) is the most common cause of epidemic gastroenteritis worldwide, and can be particularly dangerous for infants and the elderly [1,2]. Viral shedding often persists asymptomatically for weeks to months after acute infection [3,4], and is a potential source for initiation of outbreaks. Despite recent success in development of *in vitro* systems to study HNoV infection [5,6], there remains a need for robust small animal models for investigating NoV biology. Murine norovirus (MNV) shares genotypic (ssRNA, positive-sense, ~7.5kb genome) and phenotypic (fecal-oral transmission, infection of intestinal epithelial cells (IECs), persistent shedding) features with HNoV. Therefore, studies of MNV have enabled considerable advances in understanding of norovirus biology [7–9].

All noroviruses express a non-structural polyprotein encoded by ORF1 and two structural capsid proteins (VP1 and VP2) encoded by ORFs 2 and 3, respectively [10–12] (Fig. 1A). The non-structural polyprotein is cleaved by the internally-encoded viral protease into six ‘mature’ proteins (NS1/2, NS3, NS4, NS5, NS6, and NS7) [12–14]. Unlike related caliciviruses, NS1/2 of NoV remains intact for most of the virus life cycle and may be cleaved by host proteases rather than the viral protease [12,13]. NoV non-structural proteins associate with membranes and form the membranous viral replication complex [15–18]. Specific roles of NS5-7 have been defined as VpG cap, protease, and polymerase, respectively. However, the roles of NS1-4 are less well understood [19].

**Fig. 1.**
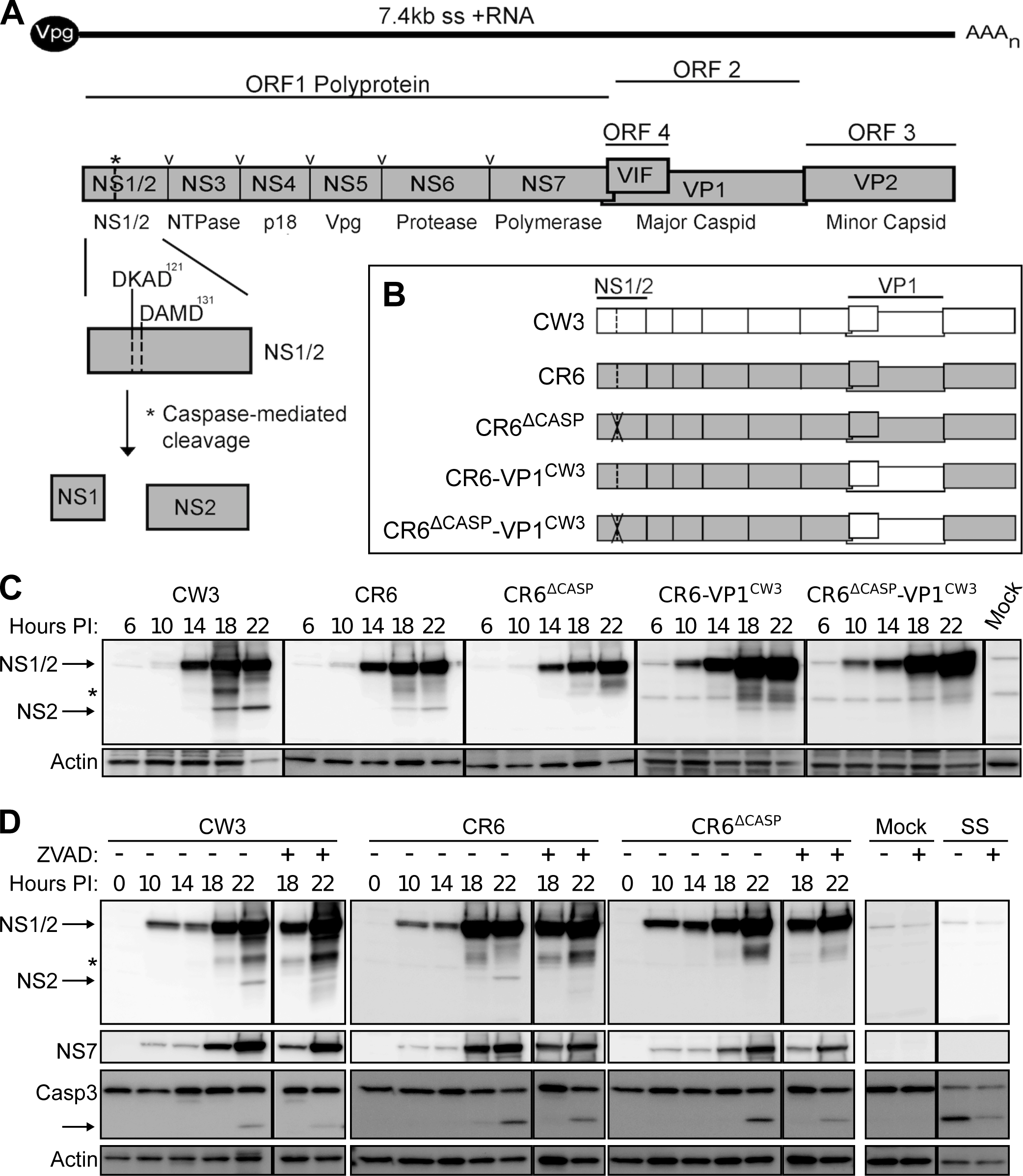
MNV NS1/2 is cleaved by host caspases at late times-post-infection. (A) Schematic of MNV genomic NS1/2 expression within the ORF1 polyprotein. Sites indicated in ORF1 are cleaved by the viral protease (**v**) and host caspases (**\***), the latter at aspartic acid residues 121 and 131 within two caspase motifs, DKAD^121^ and DAMD^131^. (B) Schematic representation of the five MNV clones used in these studies. D121G and D131G mutations are indicated by the X within NS1/2. (C) BV2 cells were infected with indicated strains of MNV (MOI=1), and whole cell extracts (WCE) were analyzed via western blot (WB) at indicated times post-infection. Image is representative of 4 independent experiments. Arrows indicate full-length NS1/2 (45kD) and NS2 (28kD). * Denotes caspase-independent bands >28kD. (D) BV2 cells were infected as in C, but subsequently treated with 50uM ZVAD-fmk at 10 hours post-infection, and WCE were probed for NS1/2, NS7, caspase 3, and actin via WB. Image is representative of 3 independent experiments. SS = staurosporine.

Our prior MNV studies compared acute strain CW3 and persistent strain CR6 to identify viral determinants of pathogenesis and persistence [20–22]. We found that the capsid gene of CW3 (VP1^CW3^) promotes extra-intestinal spread, whereas the NS1 gene of CR6 (NS1^CR6^) promotes intestinal replication and persistent fecal shedding. Persistent and non-persistent strains similarly infect macrophages, dendritic cells, T-cells, and B-cells *in vitro* and *in vivo* [6,23,24], and utilize the same cell-surface receptor, CD300lf [25,26]. However, recent studies have shown that tropism for IECs *in vivo* requires NS1^CR6^ in addition to CD300lf expression by non-hematopoietic cells [27,28]. Furthermore, only a minor lineage of IECs called tuft cells naturally express CD300lf and are susceptible to persistent infection by CR6 [27,29]. In contrast to the strain-dependent infection of IECs *in vivo*, CW3 and CR6 can replicate to similar titer in a cultured IEC cell line (M2C) that ectopically expresses CD300lf [28]. Thus, NS1^CR6^ promotes IEC persistence *in vivo* via mechanisms other than receptor usage or replicative capacity *per se*.

In addition to viral determinants of MNV infection, critical aspects of the host anti-viral response have been identified. In particular, innate control of MNV replication is determined by the compartmentalized roles of type I (α/β) and type III (λ) interferon (IFN) [30]. Both types of IFN stimulate antiviral gene expression, but the IFN-α/β receptor (IFNAR) is expressed on most cell types, whereas the IFN-λ receptor (IFNLR) is preferentially expressed on epithelial cells [30]. Correspondingly, IFN-α/β is critical for control of MNV in myeloid cells and prevents pathological dissemination [23,31,32], whereas IFN-λ is critical for control of MNV in IECs and prevents persistent shedding [33,34]. Deficiency of IFNLR promotes persistent infection, suggesting that NS1^CR6^ counteracts the IFN-λ response [28]. However, CW3 persistence is not rescued in IFNLR-deficient mice [28,34] indicating that other host pathways are also important.

In addition to the IFN response, a general host response to many diverse viral genera is initiation of apoptosis. Correspondingly, viruses have evolved ways to hijack apoptotic machinery to their benefit. Apoptosis is a non-inflammatory, highly-regulated disassembly of the cell mediated by a family of cysteine proteases (caspases). Apoptotic cellular contents are packaged and released in membranous apoptotic bodies to be taken up by professional phagocytes and other neighboring cells [35]. Because apoptosis is an innate response to infection, many viruses have developed ways to delay induction of apoptosis such as expression of anti-apoptotic Bcl-2 family homologues [36,37]. In contrast, after virus production has reached its peak, many viruses actively promote apoptotic death or utilize ‘apoptotic mimicry’ to facilitate viral dissemination [37,38]. Additionally, some viruses rely on apoptotic caspases to cleave viral proteins as an additional level of regulation [39–41]. Thus, depending on its timing in relation to viral replication, apoptosis can be anti-viral or pro-viral and is therefore regulated by a diverse array of viral mechanisms.

HNoV and MNV infection both induce apoptosis, as indicated by association with markers of apoptosis in infected intestinal tissue [42–44]. Thus, these viruses have likely also developed mechanisms to regulate apoptosis in a pro-viral manner. Apoptosis has been most extensively studied in MNV-infected myeloid cells and is associated with down-regulation of anti-apoptotic survivin protein and activation of caspases 3, 7, and 9 [45,46]. Expression of the MNV ORF1 polyprotein is sufficient to induce apoptosis [47], indicating that one or more non-structural proteins are pro-apoptotic in the absence of viral replication. Additionally, NS1/2 encodes caspase cleavage motifs that are targeted by recombinant caspase 3 in a cell-free assay and separate NS1 from membrane-associated NS2 [12]. Thus, apoptosis during MNV infection may be facilitated by non-structural proteins and host caspases may reciprocally regulate NS1/2 function [20].

Herein, we identify a critical role for NS1/2 cleavage during MNV infection and intestinal persistence. We demonstrate that at late times post-infection, concurrent with induction of apoptosis, a minority of total NS1/2 protein is cleaved in a caspase-dependent manner. Despite the relatively low proportion of total NS1/2 that is cleaved, NS1/2^CR6^ cleavage is absolutely required for persistent infection in IECs, independent of IFN-λ. Moreover, the cleavage of NS1/2^CR6^ promotes apoptotic death of infected myeloid cells and spread among cultured IECs. These data suggest that controlled induction of apoptosis during viral egress promotes spread among IECs and is required for maintenance of the persistent cellular reservoir.

## Results

### Host caspases cleave NS1/2 at late times post-infection

Previous cell-free studies identified two functional caspase motifs (DKAD^121^ and DAMD^131^) within NS1/2 that are cleaved by host caspase 3 [12] (Fig. 1A), however, the targeting of these sites during infection and functional consequences of NS1/2 cleavage was unknown. To test their role in persistent infection, we mutated these sites (D121G and D131G) within MNV strain, CR6 (CR6^Δcasp^, Fig. 1B). In parallel, we generated a variant of CR6^Δcasp^ with the capsid of the systemic strain CW3 (CR6^Δcasp^-VP1^CW3^) to enable determination of a role for NS1/2 cleavage in extra-intestinal tissues (Fig. 1B) [20,48]. Analysis of NS1/2 protein expression following infection of BV2 myeloid cells revealed a similar level of full length NS1/2 production for all strains. However, infection with CW3, CR6, or the CR6-VP1^CW3^ chimera but not NS1/2 cleavage mutant strains resulted in a ~28 kDa NS2 fragment detectable at 18-22 hours post-infection (Fig. 1C). The ~18kDa NS1 protein was inconsistently detected in cell lysate for any strain and may reflect the loss of dying cells or the small overall proportion of cleaved NS1/2. Several fragments larger than 28 kDa (marked by *) were also observed but did not consistently differ between strains, indicating they are independent of NS1/2 mutations (Fig. 1C). Overall, these data show that cleavage of NS1/2 occurs during infection, depends on caspase motifs, and is independent of other genetic variation between CW3 and CR6.

To determine the temporal relationship between caspase activity and NS1/2 cleavage, we detected cleaved caspase 3 (active form) by western blot. Cleaved caspase 3 was observed at 18-22 hours post-infection for all strains tested, concurrent with NS1/2 cleavage (Fig. 1D). To test the requirement of caspase activity for NS1/2 cleavage, we treated infected cells with the pan-caspase inhibitor, ZVAD-fmk. Inhibitor treatment reduced the proportion of cleaved caspase 3 and NS2 in cells infected with CW3 or CR6 but did not reduce the abundance of NS1/2 fragments larger than 28 kDa (Fig. 1D). Together, these data demonstrate that NS1/2 is cleaved 18-22 hours post infection with persistent or non-persistent MNV strains, concurrent with induction of apoptosis, in a caspase-dependent manner.

### NS1/2 cleavage promotes apoptosis in myeloid cells

In the course of characterizing caspase cleavage, we observed that the timing and magnitude of caspase 3 cleavage differed between viral strains (Fig. 1D). To further compare caspase activity triggered by MNV infection, we used densitometry to quantify percentage of cleaved caspase 3 and poly ADP-ribose polymerase (PARP), a known target of apoptotic caspases [49]. We observed three-fold more activated caspase 3 and 10-20% more cleaved PARP in cells 18 hours post-infection with CR6 compared to CR6^Δcasp^ or CW3 (Fig. 2A-C). There were no significant differences in MNV polymerase (NS7) detected for any infection condition suggesting that the difference in caspase 3 and PARP cleavage was not simply due to increased replication in CR6-infected cultures (Fig. 2B). Together, these data suggest that cleavage of CR6 NS1/2 (but not CW3 NS1/2) potentiates caspase activity.

**Fig. 2.**
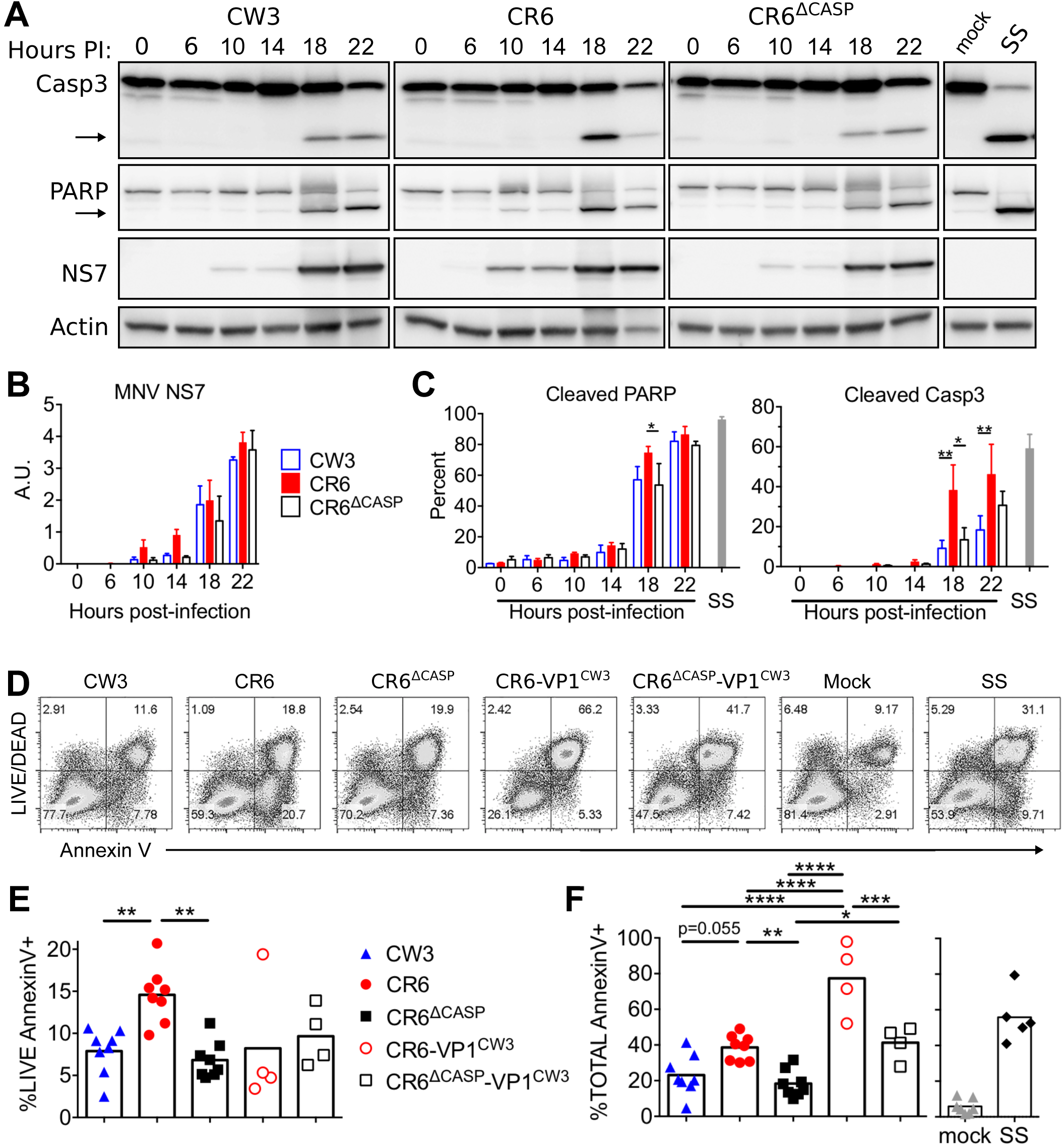
Cleavage of NS1/2 promotes apoptosis of myeloid cells. BV2 cells were infected (MOI=1) with indicated MNV strains. (A-C) Whole cell extracts were collected at indicated times and analyzed via WB for caspase 3, PARP, actin, and MNV NS7. (A) Arrows denote cleaved fragments of caspase 3 and PARP. Images are representative of three independent experiments. (B) NS7 normalized to actin analyzed via densitometry. (C) Cleaved PARP and Caspase 3 were quantified as a proportion of total protein by densitometry. (D-F) Cells were harvested at 17 hours post-infection, stained with Annexin V and LIVE/DEAD viability stain, and analyzed via flow cytometry. (D) Representative flow plots are shown from one of eight independent experiments. (E) Live, Annexin V+, and (F) total Annexin V+ cells from all experiments are graphed and mean values are indicated. Statistical significance was determined by one-way (E-F) or two-way (B-C) ANOVA, with Tukey’s multiple comparison test. *, p ≤ 0.05; **, p ≤ 0.01; ***, p≤ 0.001; ****, p≤ 0.0001; SS, staurosporine.

To test whether our panel of MNV strains differed in induction of apoptotic cell death, we quantified surface presentation of phosphatidylserine (PS, Annexin V staining) and permeability to the LIVE/DEAD viability dye by flow cytometry following infection of BV2 cells (Fig. 1B). We found that CR6 infection resulted in two-fold more apoptotic cells (Annexin V positive, LIVE/DEAD negative) than CW3 or CR6^Δcasp^ (Fig. 2A-B). In contrast to CR6, most CR6-VP1^CW3^ infected cells lost membrane integrity and were stained with the LIVE/DEAD dye (Fig. 2A-C), which is consistent with our recent findings that VP1^CW3^ drives lytic cell death and inflammatory cytokine production [48]. Even so, cells infected with CR6-VP1^CW3^ had 30-50% more total Annexin V staining than cells infected with the corresponding mutant strain CR6^Δcasp^-VP1^CW3^, indicating that NS1/2 cleavage promotes cell death in conjunction with VP1^CW3^-dependent lysis (Fig. 2C). Together, these data suggest that CR6 NS1/2 cleavage by apoptotic caspases reciprocally promotes cleavage of caspase substrates and consequential cell death.

### Cleavage of NS1/2 is not required for *in vitro* MNV replication in myeloid cells

The abundance of full-length NS1/2 and NS7 on western blots was similar between WT and mutant strains, indicating that viral protein production was unaffected by NS1/2 cleavage mutations (Fig. 1C-D, 2A). However, many additional steps beyond protein production are required for assembly of infectious virus. So, we compared production of plaque forming units (PFU) over time from myeloid cells infected with our panel of MNV strains (Fig. 1B) to assess whether production of infectious virus was dependent on CR6 NS1/2 cleavage. Mutation of NS1/2 cleavage sites did not significantly alter MNV replication kinetics or peak titers in BV2 cells following infection at a high multiplicity of infection (MOI, one PFU/cell) (Fig. 3A). Following infection of BV2 cells at a low MOI (0.01 PFU/cell), there were modest differences in early 10-18 hour titer, but all strains reached similar peak titers by 36 hours post-infection (Fig. 3A). The similar peak titers are consistent with our ability to obtain high titer viral stocks of NS1/2 cleavage mutants (generated by low MOI infection of BV2 cells) and indicate that NS1/2 cleavage is not required for MNV replication in these cells.

**Fig. 3.**
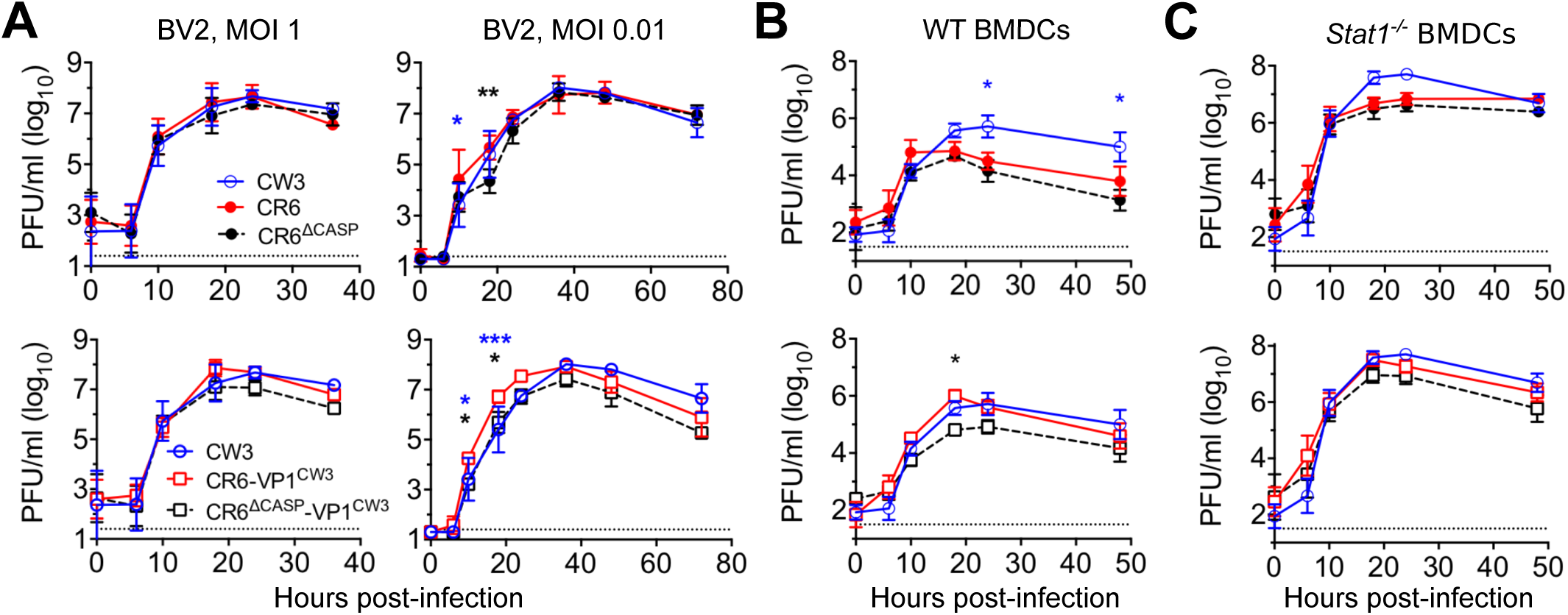
Cleavage of NS1/2 is not required for *in vitro* MNV replication in myeloid cells. BV2 cells (A), WT BMDCs (B), or *Stat1^−/−^* BMDCs (C) were infected with the indicated strain of MNV at MOI 0.01 (A) or MOI 1 (A-C) and infectious virus was analyzed at indicated times post-infection via plaque assay. Dotted lines indicate limit of detection. Error bars indicate SEM. Statistical significance was determined via two-way ANOVA with Tukey’s multiple comparisons test. *, p ≤ 0.05; **, p ≤ 0.01. BMDCs = bone marrow-derived dendritic cells. Data are an average of at least three independent experiments.

To test the requirement of NS1/2 cleavage for replication in primary myeloid cells, we generated bone-marrow dendritic cells (BMDCs) and infected them with our panel of MNV strains. Similar to BV2 cells, BMDC replication was minimally impacted by mutation of NS1/2 cleavage sites (Fig. 3B-C). However, CW3 and CR6-VP1^CW3^ reached higher peak titer in WT BMDCs compared to other strains tested, suggesting a VP1-dependent role in BMDC replication. In parallel, we tested the role of STAT1-dependent IFN responses by comparing growth of our MNV strains in *Stat1^−/−^* BMDCs. All MNV strains reached higher peak titers in *Stat1^−/−^* BMDCs compared to WT BMDCs (compare Fig. 3B and 3C), as expected from prior studies of STAT1-dependent immunity to MNV [23]. However, there were no statistically significant differences in growth that could be attributed to NS1/2 cleavage (Fig. 3C). Overall, these data indicate that NS1/2 cleavage is not required for replication of MNV in myeloid cells.

### NS1/2 cleavage is not required for lethal systemic infection

*Stat1^−/−^* mice succumb to infection in a VP1^CW3^-dependent manner due to a profound inability to control systemic replication in myeloid cells [20–22,31,32]. To test whether NS1/2 cleavage mutations impair this pathological MNV replication, we orally inoculated *Stat1^−/−^* mice with CR6-VP1^CW3^ or CR6^Δcasp^-VP1^CW3^ and monitored animals daily for signs of distress. Infection with CW3 or CR6-VP1^CW3^ resulted in 100% lethality with similar kinetics (Fig. 4A). As expected, CR6-infected controls remained healthy due to lack of VP1^CW3^-facilitated systemic spread [20]. These *in vivo* pathogenesis data suggest that NS1/2 cleavage is not required for lethal pathology resulting from uncontrolled, systemic infection of myeloid cells, consistent with no NS1/2 cleavage requirement for *in vitro* replication in myeloid cells.

**Fig. 4.**
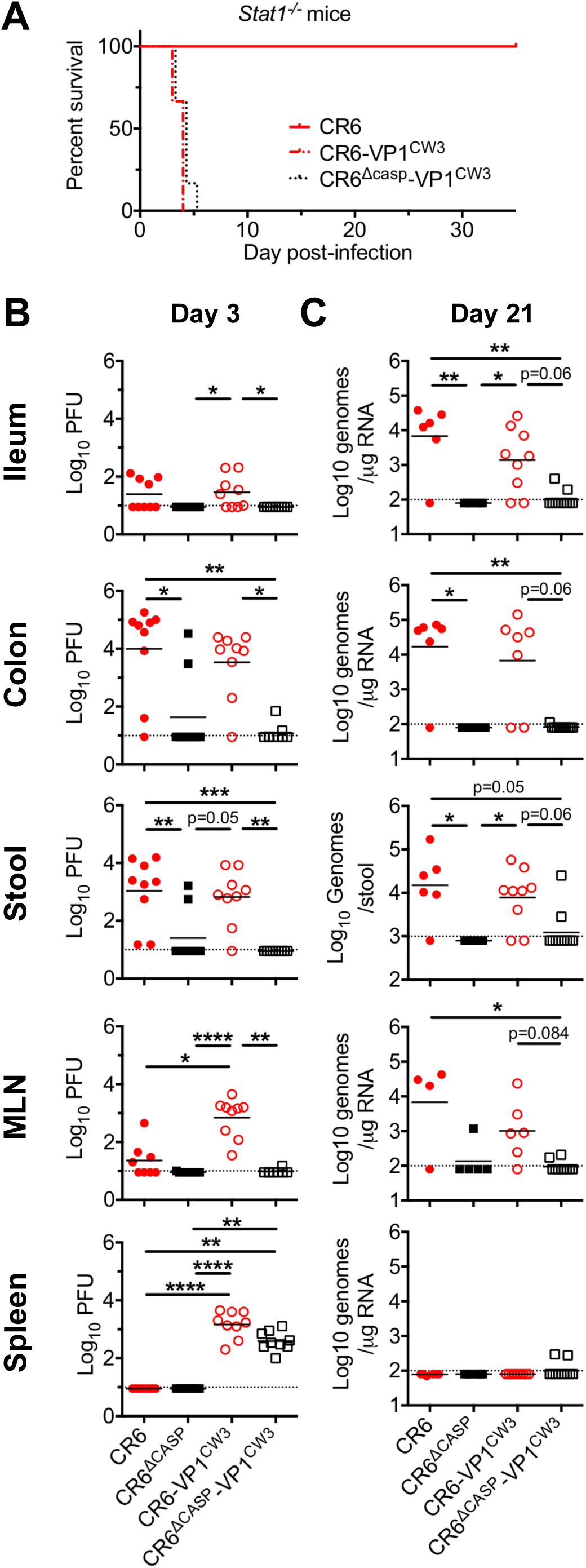
NS1/2 cleavage is required for intestinal tropism and persistence. (A) *Stat1^−/−^* mice were infected with the indicated strain and monitored daily for lethality. (B-C) WT mice were perorally infected with indicated viruses (1e6 PFU). At 3 (B) or 21 (C) days post-infection, tissues and stool were harvested and analyzed via plaque assay (B) or qPCR (D). Data are from 8-9 individual mice combined from 3 separate experiments, and mean values are indicated. Statistical significance was determined by Kruskal-Wallis test. *, p ≤ 0.05; **, p ≤ 0.01; ***, p≤ 0.001; ****, p≤ 0.0001; MLN, mesenteric lymph node.

### NS1/2 cleavage is required for intestinal tropism and persistence

To directly quantify the effect of NS1/2 cleavage on acute replication and tissue tropism, we inoculated WT mice with CR6, CR6^Δcasp^, CR6-VP1^CW3^, or CR6^Δcasp^-VP1^CW3^ and analyzed tissue and stool titers at three days post-infection (Fig. 4B). As expected from our prior studies, CR6 and CR6-VP1^CW3^ were detected on day three in the mesenteric lymph node (MLN), ileum, and colon. We also detected infectious virus shed in the feces of mice infected with CR6 and CR6-VP1^CW3^, consistent with replication in intestinal tissue. In contrast, in mice infected with NS1/2 cleavage mutant strains CR6^Δcasp^ or CR6^Δcasp^-VP1^CW3^, infectious virus was rarely detected in intestinal tissues, or being shed in the feces. However, despite severely compromised replication in ileum (10-fold) and colon (1000-fold), splenic titers of CR6^Δcasp^-VP1^CW3^ were not significantly different from CR6-VP1^CW3^ (Fig. 4B). This indicates that NS1/2 cleavage is not required for systemic spread, consistent with *Stat1^−/−^* lethality data (Fig. 4A). Overall, these acute titer data demonstrate that caspase cleavage of NS1/2 is specifically required for promoting early infection of intestinal tissue, the eventual site of persistence.

Due to limited early detection of NS1/2 cleavage mutants in the intestinal tissues, we predicted that NS1/2 cleavage would be required for intestinal persistence. Indeed, most mice infected with CR6^Δcasp^ or CR6^Δcasp^-VP1^CW3^ had undetectable viral genomes in tissues at 21 days post-infection whereas CR6 and CR6-VP1^CW3^ were present in the ileum, colon, and stool (Fig. 4C). Altogether, these findings support a role for NS1/2 cleavage in establishing early infection of the intestine and subsequent persistence.

### NS1/2 cleavage is required for intestinal persistence independent of IFN-λ

IFN-λ plays a critical role in protection against persistent MNV infection of the intestine [34], so we sought to determine whether CR6^Δcasp^ persistence was restored in the absence of IFN-λ signaling. We infected WT or *Ifnlr1^−/−^* mice with CR6 or CR6^Δcasp^ and monitored shedding in the stool over time (Fig. 5A, B), and persistence in the colon at 21 days post-infection (Fig. 5C, D). Additionally, to determine whether an early barrier to MNV infection could be overcome with increased infectious dose, we included an experimental group inoculated with a 10-fold higher dose of CR6^Δcasp^ (1e7 PFU). MNV was shed in the stool of all WT mice infected with CR6 beginning on day three and continuing until the experimental endpoint, day 21. Consistent with the known role of IFN-λ, *Ifnlr1^−/−^* mice infected with CR6 shed greater than 10-fold more virus relative to WT mice (Fig. 5A, B), and CR6 was detected in colon tissue at levels commensurate with fecal shedding (Fig. 5C, D). In contrast, we were unable to detect shedding or colonic persistence of CR6^Δcasp^ at any time from WT or *Ifnlr1^−/−^* mice (Fig. 5A, B). These data indicate that CR6 NS1/2 cleavage promotes persistent infection via a novel mechanism separate from IFN antagonism.

**Fig. 5.**
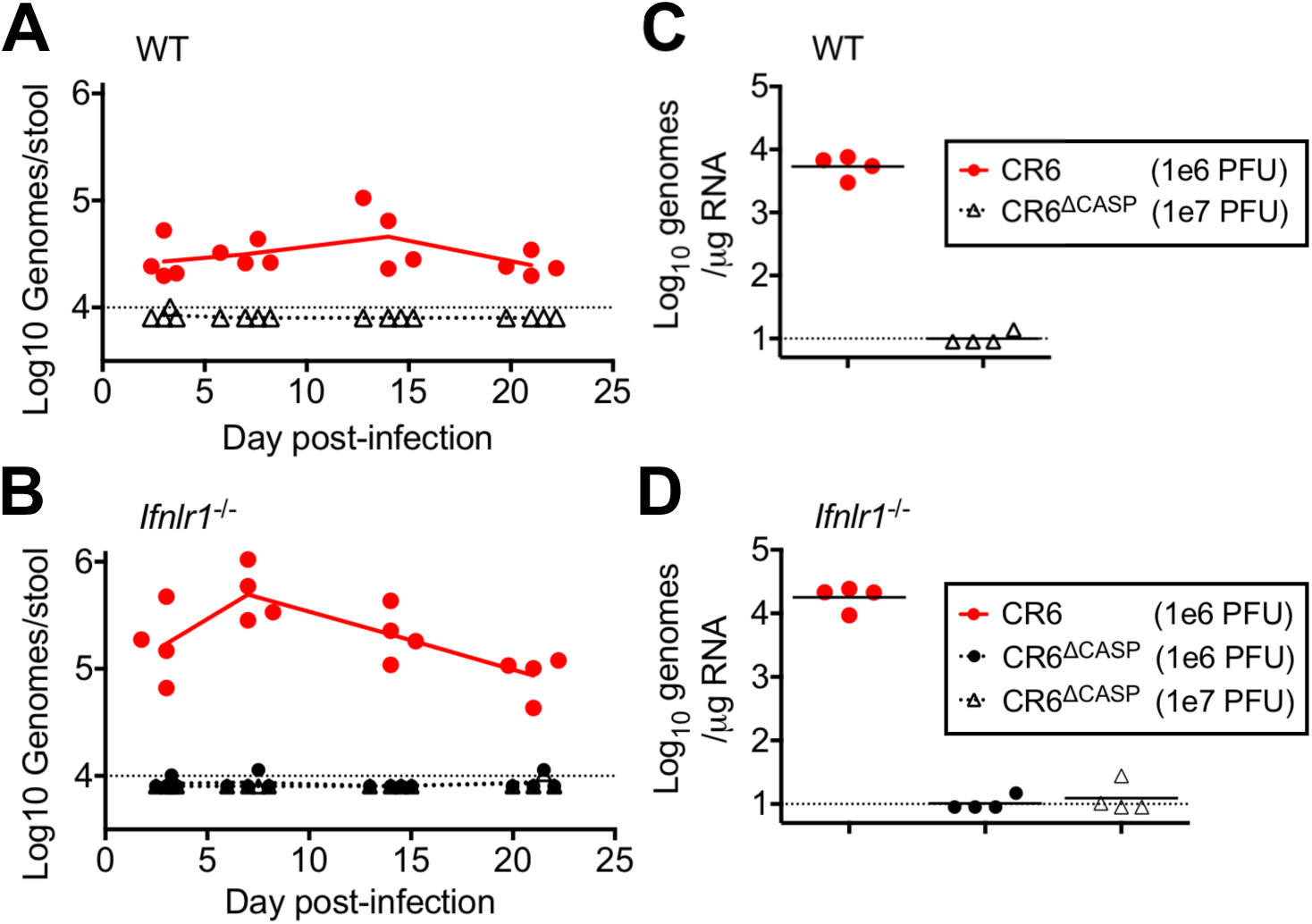
Caspase cleavage of NS1/2 is required independent of the IFN-λ response. WT (A, C) or *Ifnlr1^−/−^* (B, D) mice were infected by oral gavage with either 1e6 PFU or 1e7 PFU, as indicated. Fecal samples (A, B) were collected at 3, 7, 14, and 21 days post-infection, and colon tissue (C, D) was harvested at 21 days post-infection. All samples were analyzed via RT-qPCR, and the mean of each group (N=4) is indicated. Dotted lines indicate limit of detection.

### Cleavage of NS1/2 is critical for MNV infection of intestinal epithelial cells

Our preceding data demonstrated that disrupting NS1/2 cleavage prevents intestinal infection and persistence in either WT or *Ifnlr1^−/−^* mice (Fig. 4-5). Because intestinal persistence requires establishment of an infected IEC reservoir, we hypothesized that NS1/2 cleavage promoted early IEC tropism. To test this hypothesis, we infected WT or *Ifnlr1^−/−^* mice with our panel of MNV strains (Fig. 1B) and quantified IEC infection at three days post-infection by flow cytometry. We pooled IECs isolated from proximal colon, cecum, and distal small intestine (devoid of Peyer’s patches), and stained intracellularly for MNV NS1/2 (Fig. 6A-C). We rarely detected infected IECs in WT mice at this early timepoint, indicating that acute infection of IECs in our WT mice is below the limit of detection for this assay (Fig. 6B). However, IEC infection was increased in *Ifnlr1^−/−^* mice relative to WT mice, indicating that IFN-λ limits early establishment of infection in the IEC reservoir (Fig. 6A, C). CR6 and CR6-VP1^CW3^ were detected in an average of 40 IECs per million in *Ifnlr1^−/−^* mice, but we did not detect significant infection of IECs by NS1/2 cleavage mutants (Fig. 6C). To contextualize IEC data, we quantified MNV infection of non-epithelial cells in Peyer’s patches by qPCR. We detected viral RNA in Peyer’s patches for the majority of CR6-infected mice, ~50% of CR6^Δcasp^-infected mice, and all mice infected with CR6-VP1^CW3^ or CR6^Δcasp^-VP1^CW3^ (Fig. 6D, E). These data suggest that NS1/2 cleavage is not strictly required for initial infection of non-epithelial cells during acute infection, particularly for the CR6-VP1^CW3^ strain that robustly infects Peyer’s patches and spleen (Figs 4B, 6C, D). Furthermore, the relationship between replication of MNV strains in Peyer’s patches of *Ifnlr1^−/−^* mice was similar to WT mice (Fig. 6C, D), consistent with a minimal role of IFN-λ in non-epithelial cells. Together, these data indicate that NS1/2 cleavage promotes early infection of IECs, independent of early replication in Peyer’s patches.

**Fig. 6.**
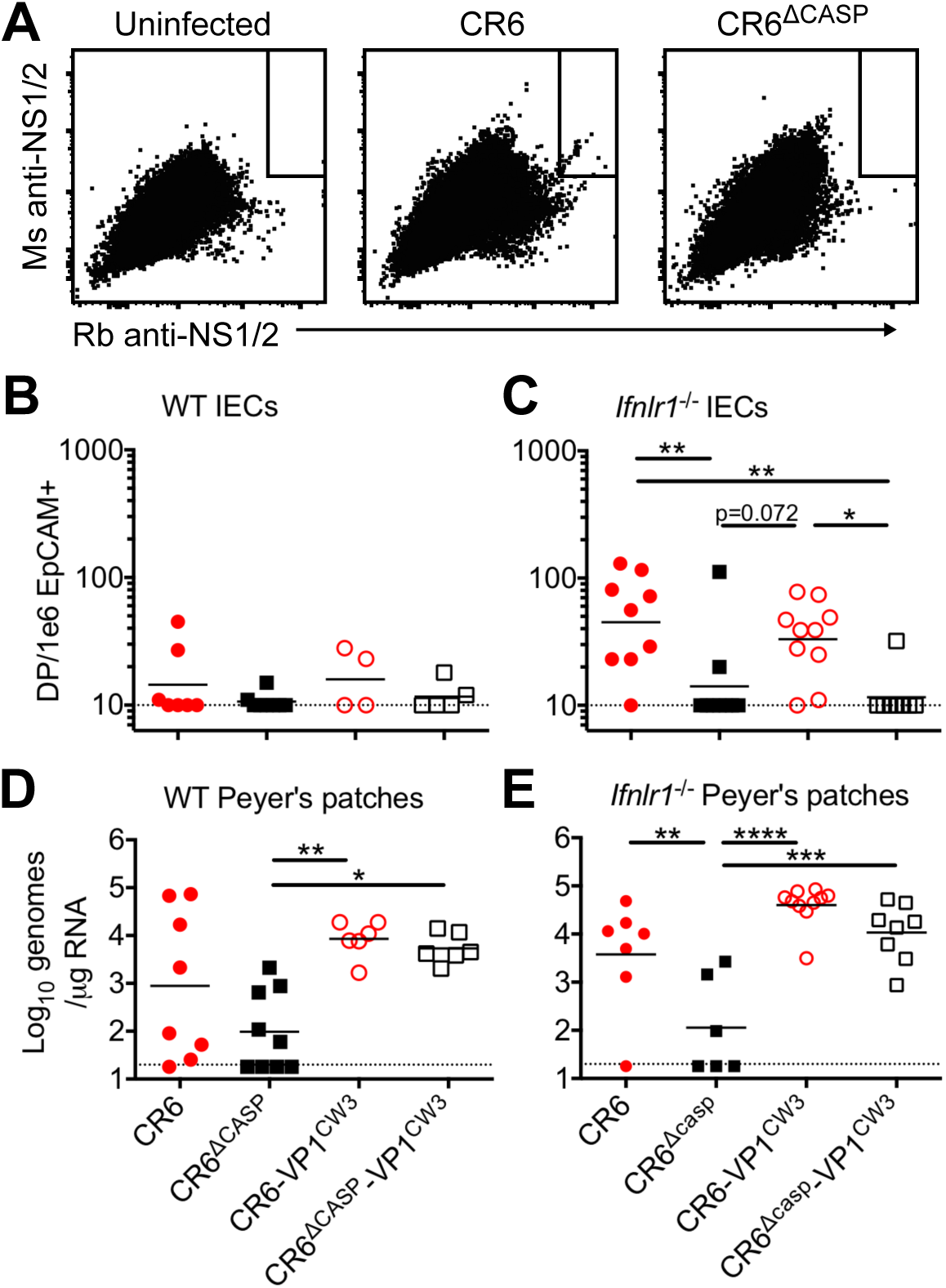
NS1/2 cleavage is necessary for infection of intestinal epithelial cells. Mice deficient for the IFN-λ receptor (A, C, E) or matched receptor-sufficient controls (B, D) were infected with indicated viruses and analyzed at three days post-infection. (A-C) Intestinal epithelial cells (CD45-, EpCAM+) were collected and doubly stained for NS1/2 as described in methods. (A) Flow plots representing concatenated data from five mice in C. Dotted line indicates limit of detection. (D-E) Peyer’s patches were collected and analyzed via qPCR for MNV genomic RNA and normalized to RPS29. Data are combined from at least three separate experiments. Statistical significance determined by one-way ANOVA with Tukey’s multiple comparisons test. *, p ≤ 0.05; **, p ≤ 0.01; ***, p≤ 0.001; ****, p≤ 0.0001.

In complementary experiments, we infected mice with our panel of MNV strains and detected MNV genomes and EpCAM transcripts in small intestine sections via RNAscope at 48-72 hours post-infection (Fig. 7). We included groups of WT and *Ifnar1 × Ifnlr1* double knockout (DKO) mice to compare IFN-restricted and unrestricted tropism, respectively. We reproducibly detected CR6 infection of IECs in WT mice at an average of 1 double positive (MNV and EpCAM) cell per cm of tissue length, whereas CR6-VP1^CW3^ was not consistently detected in IECs of WT mice and NS1/2 cleavage mutants were not detected at all (Fig. 7A-B). In DKO mice, CR6 and CR6-VP1^CW3^ were detected with increased frequency relative to WT mice at seven and one DP cells per cm tissue length, respectively (Fig. 7C-D). In contrast, infection of IECs by NS1/2 cleavage mutants remained below the limit of detection in DKO mice (Fig. 7E). Infection of non-epithelial cells was observed inconsistently for all strains in WT mice and did not correlate with IEC infection; in DKO mice, CR6-VP1^CW3^ and CR6^Δcasp^-VP1^CW3^ robustly infected non-epithelial cells, whereas CR6 infection of non-epithelial cells remained inconsistent (Fig. 7F). Taken together, these data demonstrate an IFN-independent role of NS1/2 cleavage in promoting early infection of IECs, the cellular reservoir of MNV persistence.

**Fig. 7.**
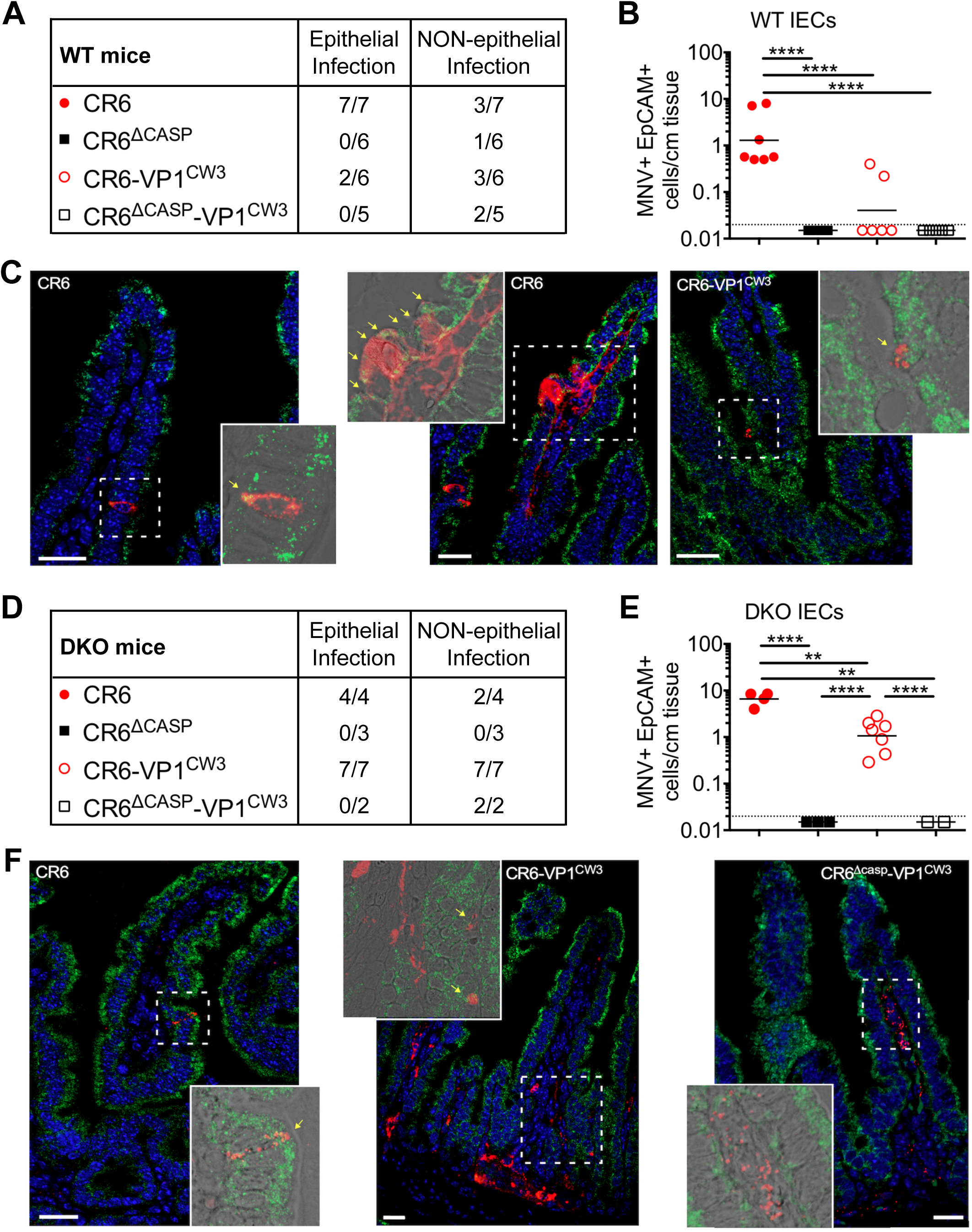
NS1/2 cleavage is necessary for infection of intestinal epithelial cells. Ileum tissue from MNV-infected WT (A-C) or DKO (D-F) mice was analyzed via RNAscope at 2-3 days post-infection. MNV RNA is in red, EpCAM is in green, and nuclei are stained with DAPI in blue. The tables in A and D indicate the total number of mice analyzed and the number of mice with MNV genome-positive cells in epithelial (EpCAM-positive) or non-epithelial (EpCAM-negative) compartments. Yellow arrows in images indicate MNV+ EpCAM+ cells. Data points in B and E represent individual mice and indicate the number of MNV+ EpCAM+ cells per cm of tissue. White scale bar, 20µm. Statistical significance determined by one-way ANOVA with Tukey’s multiple comparisons test. **, p ≤ 0.01; ****, p≤ 0.0001.

### NS1/2 cleavage promotes spread among IEC monolayers

The infected IECs we observed *in vivo* on day three may be a result of viral amplification and spread rather than initial infection. To directly test the role of NS1/2 cleavage on initial infection of IECs, we used a previously characterized CD300lf-transduced IEC cell line (M2C-CD300lf, hereafter M2C), which can be infected by CW3 and CR6 [28]. We infected M2C cells with our panel of persistent and non-persistent strains (Fig. 1B) and quantified replication over time by plaque assay (Fig. 8A). Following infection at an MOI of one, persistent strains (CR6 and CR6-VP1^CW3^) produced significantly more infectious progeny at 10 hours post-infection compared to non-persistent strains (CW3, CR6^Δcasp^, or CR6^Δcasp^-VP1^CW3^). However, between 18-48 hours post-infection all strains reached similar peak titers with at most a two-fold difference between persistent and non-persistent strains (Fig. 8A). Likewise, infection of M2C cells at a low MOI (0.01 PFU/cell) resulted in higher titers for persistent strains at 10 hours post-infection, but all strains reached similar titers by 18 hours post-infection (Fig. 8B). However, between 24-72 hours following low-MOI infection, persistent strains proceeded to replicate robustly and reach similar peak titer as high MOI infection, whereas non-persistent strains replicated more slowly with 10-fold lower titer between 24-72 hours (Fig. 8B). These findings differ from our earlier analysis in myeloid cells, where higher peak titers did not correlate with persistence phenotype (Fig. 3), and suggest specific roles of CR6 NS1/2 cleavage in IEC infection.

**Fig. 8.**
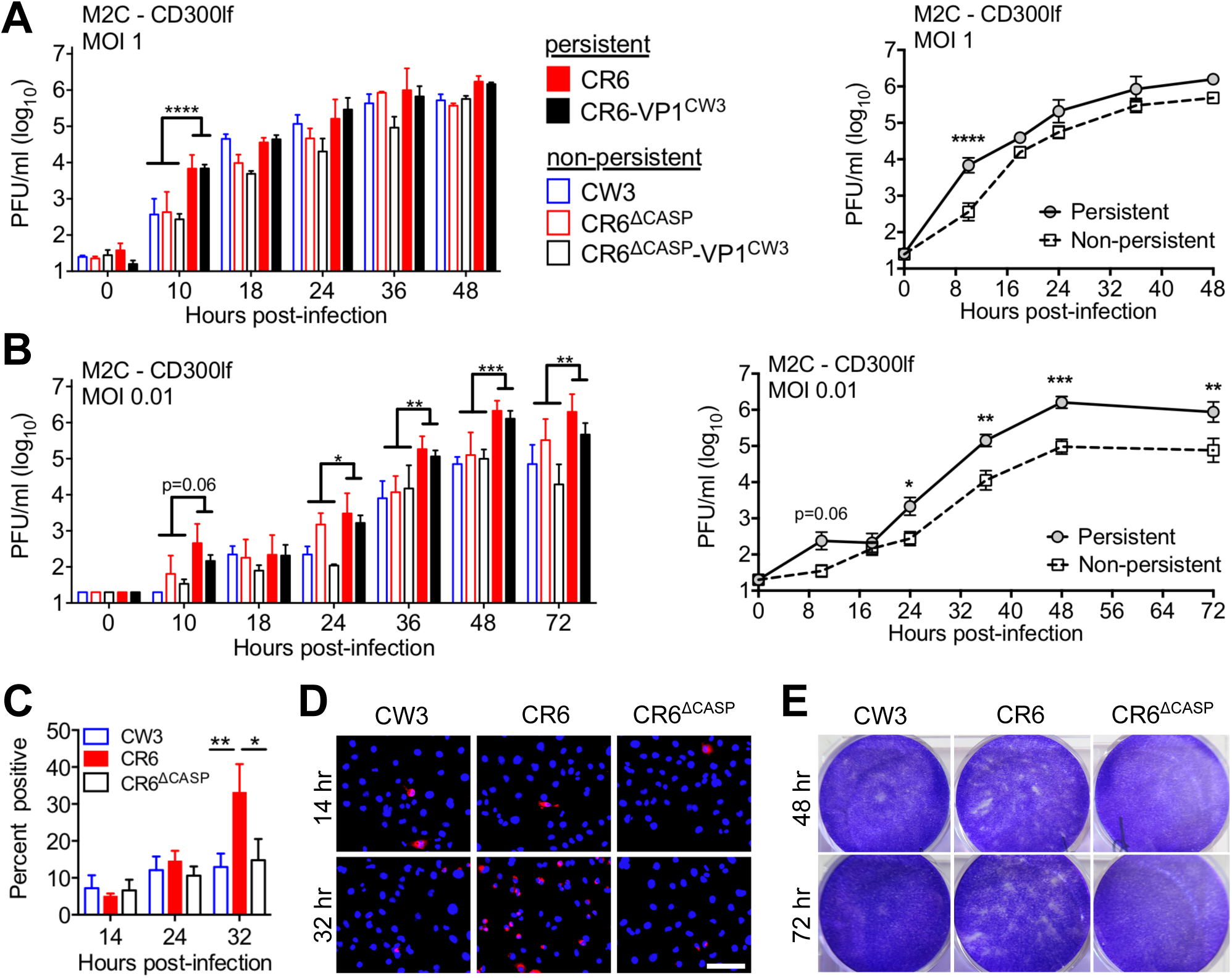
NS1/2 cleavage promotes spread among IEC monolayers. (A-B) M2C-CD300lf cells were infected with the indicated strains at MOI of 1 (A) or 0.01 (B) and production of infectious MNV was monitored over time by plaque assay on BV2 cells. Curves show growth of strains grouped by persistence phenotype. (C-E) M2C-CD300lf cells were infected with the indicated strains at MOI of 1 (C-D) or 0.1 (E) and stained for MNV NS7 immunofluorescence microscopy at the indicated time post-infection (C-D) or stained with crystal violet dye to visualize the cell monolayer (E). Data in A-C is grouped from at least three independent experiments. D and E are representative images, and white scale bar is 100µm. Statistical significance determined by two-way ANOVA of grouped data with Tukey’s multiple comparison test. *, p ≤ 0.05; **, p ≤ 0.01; ***, p≤ 0.001; ****, p≤ 0.0001.

To further characterize infection in IECs, we quantified infected cells by immunofluorescence staining for MNV NS7 following infection at MOI of one. Unexpectedly, only 10% of M2C cells were NS7-positive at 14 and 24 hours post-infection, suggesting that MNV does not efficiently initiate replication in these cells. However, strains CW3, CR6, or CR6^Δcasp^ all had a similarly low percentage of initially infected cells (Fig. 8C, D). This equal infection suggests that early differences in titer at 10 hours are due to different rates of virus production per cell. Although the initial proportion of infected cells was similar between strains, the percentage of CR6-infected cells increased from 10 to 30% by 32 hours whereas the percentage of CW3 or CR6^Δcasp^ infected cells did not significantly increase (Fig. 8C, D). Furthermore, visual inspection of CR6-infected M2C cell monolayers at 48-72 hours post-infection revealed patches of minimal cellularity whereas CW3 or CR6^Δcasp^ infected monolayers remained intact (Fig. 8E). In sum, these data indicate that non-persistent strains are similarly able to initially infect IECs, but spread among IEC monolayers is facilitated by cleavage of CR6 NS1/2.

### NS1/2 cleavage promotes death of IECs

We next sought to identify the cellular response altered by NS1/2 cleavage that may account for differential spread among IEC monolayers and establishment of IEC infection *in vivo*. Analyses in myeloid cells indicated that NS1/2 cleavage promoted apoptotic cell death (Fig. 2). To test whether IEC apoptosis was also promoted by NS1/2 cleavage, we infected M2C monolayers with CW3, CR6, or CR6^Δcasp^ and characterized cleavage of NS1/2, caspase 3, and PARP by western blot. In contrast to the robust caspase 3 cleavage seen in infected BV2 cells, we detected less than 10% cleavage of caspase 3 in M2C lysates from infection with any strain at any timepoint and were infrequently able to detect NS1/2 cleavage at any time post-infection (Fig. 9A). This reduced cleavage is consistent with the relatively low frequency of infected M2C cells and loss of cellularity in CR6-infected cultures (Fig. 8C-E). Additionally, staurosporine treatment of M2C resulted in visible loss of cellularity, but less cleavage of caspase 3 (25% vs. 60%) and PARP (10% vs. 95%) compared to BV2 cells (Fig. 9A-E, 2A-E). Despite the relatively low amount of caspase cleavage, M2C monolayers infected with CR6 had three-fold more caspase 3 and PARP cleavage than those infected with CW3 or CR6^Δcasp^ (Fig. 9A-C) with no significant difference in viral protein production (NS7, Fig. 9A, B). To determine if differences in caspase activity of infected M2C cells correlated with a different proportion of apoptotic cells, we quantified Annexin V staining and cell lysis by flow cytometry. No apoptotic cells were detected in the first 24 hours post-infection (not shown), consistent with slower replication kinetics and caspase activation (Fig. 8, 9A-C). However, by 36 hours post-infection CR6 triggered a three-fold increase in apoptotic cells (Annexin V positive, LIVE/DEAD negative) whereas CW3 or CR6^Δcasp^ triggered only a two-fold increase over background (Fig. 9D, E). Altogether, these data demonstrate that CR6 NS1/2 cleavage promotes apoptotic caspase activity and cell death of myeloid (BV2) and IEC (M2C) cell lines.

**Fig. 9.**
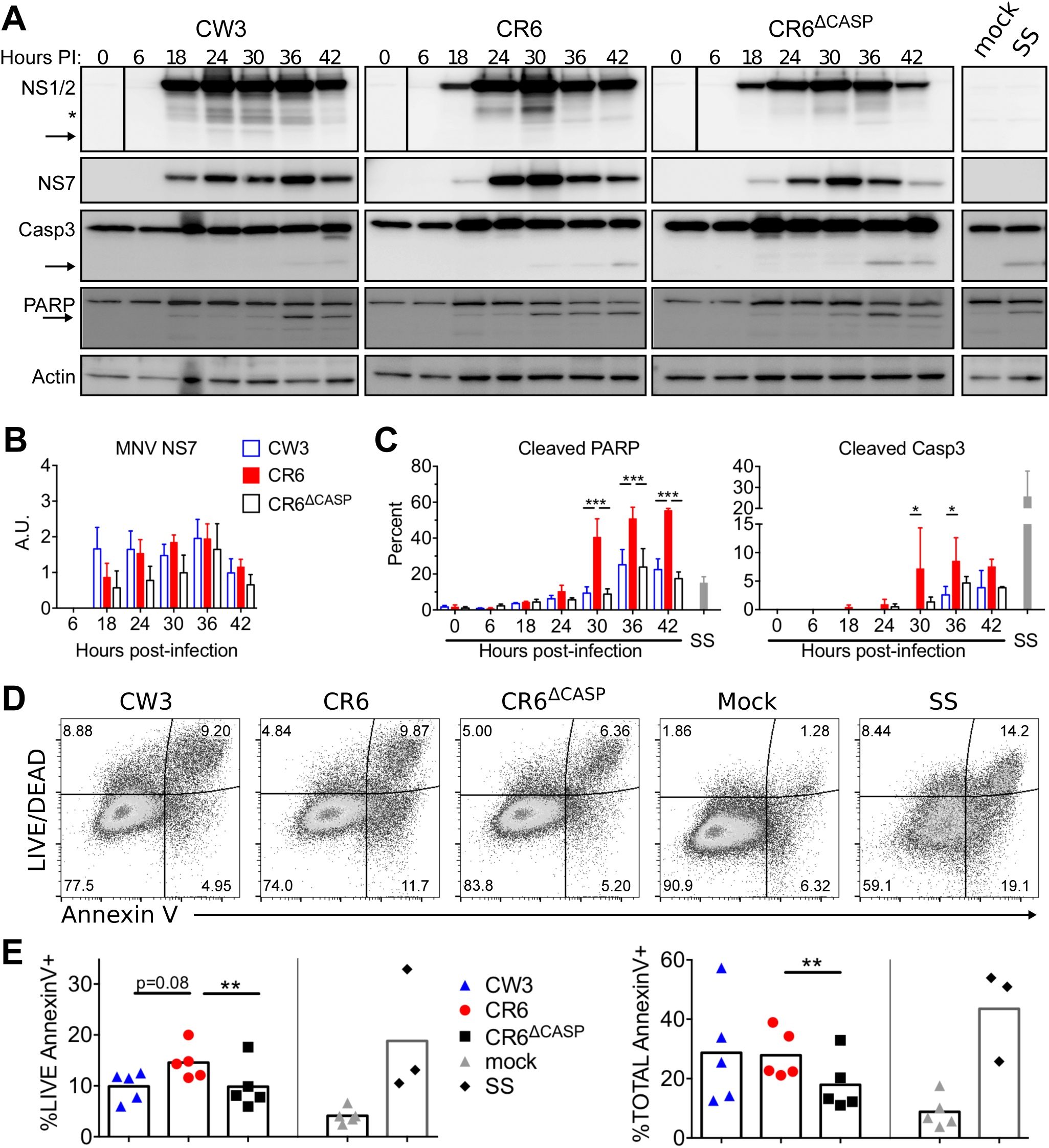
NS1/2 cleavage promotes death of IECs. M2C-CD300lf cells were infected (MOI=1) with indicated MNV strains. (A-C) Whole cell extracts were collected at indicated times and analyzed via WB for MNV NS1/2, caspase 3, PARP, actin, and MNV NS7. (A) Arrows denote cleaved fragments of NS1/2, caspase 3 and PARP. Images are representative of three independent experiments. (B) NS7 normalized to actin analyzed via densitometry. (C) Cleaved PARP and Caspase 3 were quantitated as a proportion of total protein by densitometry. (D-E) Cells were harvested at 36 hours post-infection, stained with Annexin V and LIVE/DEAD viability stain, and analyzed via flow cytometry. (D) Representative flow plots. (E) Live, Annexin V+, and total Annexin V+ cells are graphed and mean values are indicated. Data is from three to five independent experiments. Statistical significance was determined by one-way (E) or two-way (B-C) ANOVA with Tukey’s multiple comparison test. *, p ≤ 0.05; **, p ≤ 0.01; ***, p ≤ 0.001; SS, staurosporine.

MNV-induced death occurred in a relatively low proportion of M2C cells over an extended time course with strain-dependent loss of cellularity (Fig. 8E). To more comprehensively compare strain-dependent differences in this context, we used the incucyte imaging system to quantify confluency, Annexin V staining, and cytotox viability dye staining over time. Control M2C monolayers treated with staurosporine died rapidly with loss of confluency and increasing Annexin V and cytotox staining between 4-12 hours post-treatment (Fig. 10). CR6 infected monolayers also lost confluency over the course of the experiment whereas CW3 or CR6^Δcasp^ infected monolayers remained intact (Fig. 10A). Monolayers infected with all MNV strains became positive for Annexin V and cytotox with similar kinetics, beginning at 24 hours post-infection and increased to a maximum at 48 hours (Fig. 9A, B, S1-S5). However, CR6 infection resulted in more overall Annexin V and cytotox staining over the course of the experiment than CW3 or CR6^Δcasp^ infection, as quantified by area-under-curve (AUC) analyses (Fig. 9C). Although flow cytometry experiments detected Annexin V positive M2C cells with intact membranes (Fig. 9D, E), these live imaging data suggest that there is a rapid progression from apoptosis to lytic death in virally infected cells. Overall, these analyses of cell death suggest that CR6 NS1/2 cleavage promotes increased apoptotic and lytic death in IECs. Taken together with our *in vivo* data, we propose that NS1/2 is a viral sensor of caspase activity whose cleavage promotes execution of cell death programs during viral egress, lateral spread among intestinal epithelial cells, and persistent shedding (Fig. 11).

**Fig. 10.**
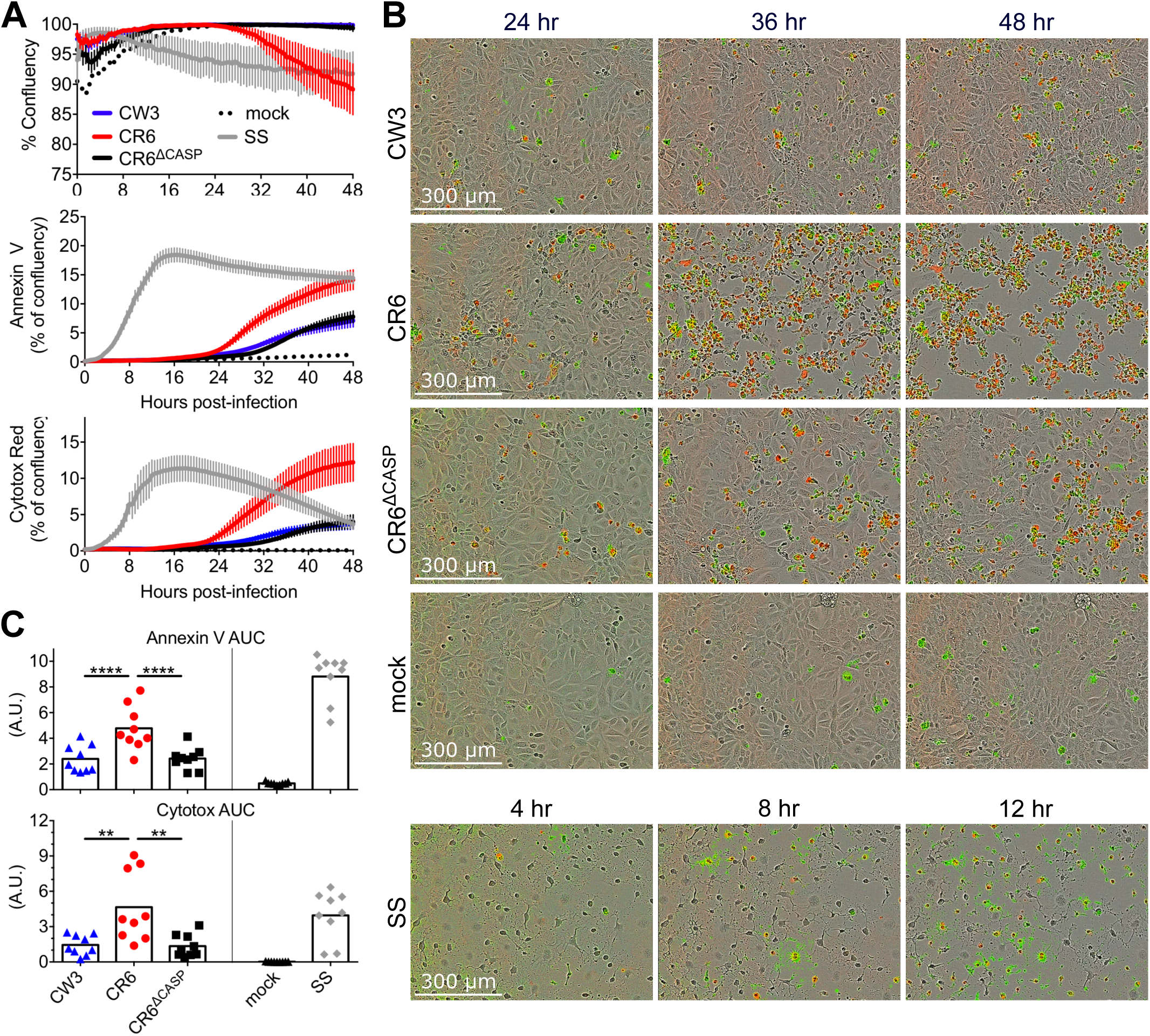
NS1/2 cleavage promotes death of IECs. M2C-CD300lf cells were infected (MOI=1) with indicated MNV strains. Cells were imaged for 48 hours on the incucyte platform in the presence of Annexin V and cytotox red. (A) Quantitation of confluency, Annexin V area normalized to confluency, and cytotox red area normalized to confluency over time. (B) Representative images from indicated times post-infection. See also videos S1-S5. (C) Area under the curve was quantitated for data in A. Data points indicate individual wells. Data in A and C is combined from three independent experiments. Statistical significance in C determined by one-way ANOVA with Tukey’s multiple comparison test. **, p ≤ 0.01; ****, p ≤ 0.0001; AUC, area under curve; SS, staurosporine.

**Fig. 11.**
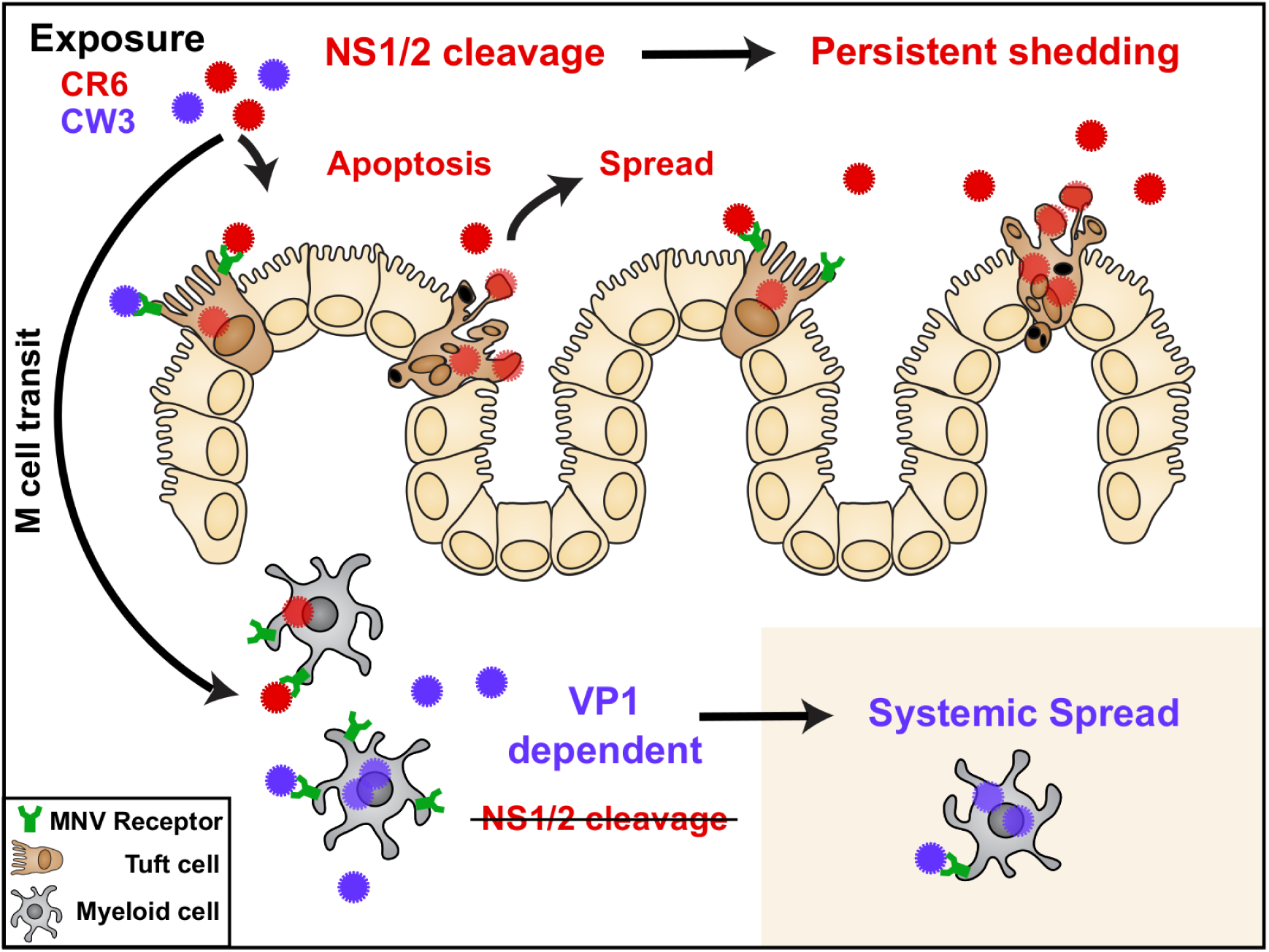
Model. MNV can cross the epithelial barrier via M cells and replicate in myeloid cells therein [24,67]. The capsid (VP1) of non-persistent strain CW3 (blue) promotes acute replication in intestinal myeloid cells and is required for spread to myeloid cells of systemic tissues. A different MNV gene, NS1 of persistent strain CR6 (red) is required for establishment of infection in rare CD300lf-expressing IECs (tuft cells), the cellular reservoir of persistent intestinal infection and fecal shedding. Cleavage of CR6 NS1 from precursor NS1/2 promotes robust tuft cell infection by potentiating apoptotic caspase activity, death of infected tuft cells, and spread to dispersed target cells within the intestinal epithelium. The role for NS1 and NS1/2 cleavage is specific to the intestinal epithelium because NS1/2 cleavage is not required for systemic replication.

## Discussion

Herein, we demonstrate that caspase-mediated cleavage of the nonstructural protein, NS1/2, is critical for persistent MNV infection of the intestine. We engineered and characterized a strain of MNV in which NS1/2 is not cleaved (CR6^Δcasp^), and our *in vitro* and *in vivo* analyses demonstrate that NS1/2 cleavage is not required for viral replication in myeloid cells but is specifically required for promoting infection in IECs, the cellular reservoir of persistence. Furthermore, our data indicate that NS1/2 cleavage potentiates apoptosis and lytic death, suggesting that regulation of cell death is a critical aspect of establishing a persistent MNV infection in IECs.

NS1/2 was not previously known to play a role in cell death during MNV infection, but is required for viral replication beginning at the earliest stages of replication complex assembly. Norovirus replication occurs in the cytoplasm on cellular membranes with a variety of organelle markers [15], and occurs near membranous vesicles [15,23]. MNV NS1/2 localizes to the endoplasmic reticulum (ER) [16,17] and promotes replication via membrane re-organization and through an interaction with the vesicle-associated membrane protein, VAPA [18,50]. Similarly, HNoV NS1/2 interacts with VAPA, localizes to vesicles, and disrupts vesicle trafficking when ectopically expressed [18]. Our data indicate that this early and essential role of NS1/2 remains intact when the caspase cleavage sites are mutated, and viral replication occurs normally in myeloid cells (Fig. 2). NS1/2 cleavage only occurs after viral replication has peaked indicating that its role in promoting death, IEC infection, and persistence involves relatively late events in the viral life cycle.

The last step of the MNV replication cycle is egress, which occurs concurrently with induction of apoptosis. Therefore, NS1/2 cleavage and associated apoptotic death may be important for regulating release of viral progeny. How noroviruses are released from infected cells is not well understood, but infection of macrophages, DCs, or IECs does result in cell lysis ([23], Figs. 2, 9, 10) and non-enveloped viruses generally rely on cell lysis to release newly assembled virions. Apoptotic processes may alter egress by reducing lytic virion release and favoring retention of virions within cellular membranes. Indeed, a number of non-enveloped viruses utilize vesicle-associated egress as a means of evasion from immune detection, and vesicles with exposed phosphatidylserine allow for phagocytic uptake of novel target cells that may not express the viral receptor [51,52]. In addition to facilitating phagocytic uptake, membranes with PS can inhibit IFN responses via activation of Tyro3/Axl/Mer (TAM) receptor tyrosine kinases on target cells [53], which would allow more efficient viral replication. Thus, vesicular egress has the potential to dramatically change the outcome of infection via biophysical and immunological mechanisms. Intriguingly, recent studies by the Altan-Bonet group indicate that MNV can be released from intact cells in PS-positive membranes [54] but the proportion of total virions released in this manner remains unknown. Therefore, it will be important in future work to consider the role of NS1/2 cleavage on regulation of this newly-described membranous egress and the role of viral association with membranes on spread among IECs.

The MNV receptor CD300lf is specifically expressed on MNV target cells, including myeloid cells and a specialized IEC subtype, tuft cells [25]. Tuft cells are a rare cell type in the intestinal epithelium and are dispersed among non-tuft, CD300lf-negative IECs [55]. Our analyses often found intestinal villi with multiple, adjacent epithelial cells infected (Fig. 7C), raising the possibility of direct MNV spread to neighboring IECs, which could be mediated by cell death or apoptotic egress and regulated by NS1/2 cleavage. Our *in vitro* studies of infection in IEC monolayers support this hypothesis, and the dispersed nature of CD300lf-expressing tuft cells poses a strong selective pressure for efficient spread across receptor-negative stretches of the intestinal epithelium (Fig. 11). Recent development of *in vitro* methods for differentiating tuft cells in primary IEC cultures [56] will enable future experiments to characterize spread in an IEC monolayer more representative of the *in vivo* scenario.

Regular turnover of the intestinal epithelium results in a short lifespan for differentiated epithelial cells, which undergo a specialized form of apoptotic cell death (anoikis) triggered as the cell detaches from the basal membrane and is shed into the lumen [57]. Specific Bcl-2 family proteins are critical players in anoikis [58,59], and display increased expression as IECs transit up the villus [59]. It is possible cleavage of NS1/2 (and perhaps NS1 directly) interferes with Bcl-2 family proteins as a means of altering the kinetics of the highly-regulated pathways that control apoptosis within differentiated IECs. Interestingly, similar to NS1/2 cleavage promoting apoptosis during infection, some Bcl-2 proteins (e.g. Bid) are also regulated via cleavage, in which the truncated version is the active, pro-apoptotic form of the protein [60]. An important future direction will be to examine whether NS1/2 interactions with the apoptotic machinery plays a role in maintaining IEC infection.

As mentioned above, PS receptor activation associated with apoptotic death could limit host responses. In contrast, virus production that results from cell lysis would promote inflammatory signals to alert the host of an infection. In addition to professional phagocytes, phagocytosis by non-professional phagocytes (e.g. epithelial cells) is critical for efficient clearance of apoptotic cells, and for limiting inflammation in tissues with high cellular turnover [61]. Specifically, in a mouse model of colonic injury, phagocytic uptake of apoptotic cells by IECs was sufficient for limiting inflammation in the colon [62], and phagocytosis mediated by epithelial cells is similarly critical for reducing inflammation in lung tissue [63] and mammary glands [64]. Thus, intestinal infection that promotes apoptotic death and the phagocytic uptake by IECs would limit host detection and provide a non-inflammatory environment that would allow more efficient infection of IECs and benefit MNV persistence. Future work will seek to identify the contribution of these immunoregulatory pathways and apoptotic viral egress on IEC infection. Our discovery of NS1/2 as an “apoptosis sensor” that reciprocally regulates apoptotic death and facilitates epithelial spread (Fig. 11) has opened the door to such mechanistic studies and expanded our understanding of the requirements for viral persistence in the intestine.

## Materials and Methods

### Cells, virus, and plaque assays

BV2 cells (gift from Herbert Virgin IV, Washington University, St. Louis, MO), a mouse microglial cell line [65], were maintained in DMEM (Life Technologies, ThermoFisher, Carslbad, CA) with 5% fetal bovine serum (FBS) (VWR Seradigm, Radnor, PA), 1X Penicillin/Streptomycin/Glutamine (P/S/G) solution, and 10mM HEPES. M2C-CD300lf cells, a murine colonic epithelial cell line transduced with the MNV receptor, CD300lf [28], were maintained in DMEM with 10% FBS, 1X P/S/G, and 10mM HEPES. For bone-marrow derived dendritic cells (BMDCs), bone marrow was isolated from the long bones of each hind leg of either a WT or *Stat1^−/−^* mouse and suspended in RPMI (Life Technologies). Red blood cells were lysed using red blood cell lysis solution (Sigma, St Louis, MO) and remaining bone marrow cells were cultured in RPMI with 10% FBS, 1X P/S/G solution, 10mM HEPES, 1X Sodium Pyruvate, 1X non-essential amino acids, and 20ng/mL GM-CSF obtained from supernatants of GM-CSF-secreting cells (J558L, [66]). After seven days of culture (37C, 5% CO_2_), non-adherent cells were collected, infected in suspension with indicated MNV strains (MOI = 1), and plated without GM-CSF for the indicated time. For indicated experiments, ZVAD-fmk (Cell Signaling Technology) was re-suspended in DMSO and added to cells at 70 μM. As a positive control for apoptosis assays, cells were treated with 1uM staurosporine (SS) (CST) for 3.5h, at 37C.

CR6^Δcasp^ MNV was generated using site directed mutagenesis as previously described [20] with the following primers: CR6 D_121_G, 5’GCCTAAGGAAGATAAAGCC<u>G**G**T</u>GCGCCCTCCCATGCG and CR6 D_131_G, 5’-TGCGGAGGACGCCATG<u>G**G**T</u>GCAAGGGAGCCCATAATTGG. The targeted residue is underlined with the mutation in bold. Correct mutations were verified via sanger sequence analysis.

MNV stocks were generated from plasmids, as described [22]. Briefly, plasmids encoding viral genomes of parental CW3 (GenBank accession no. <u>EF014462.1</u>), parental CR6 (GenBank accession no. <u>JQ237823</u>), CR6-VP1^CW3^ [20,22] and the two NS1/2_Δcasp_ viral genomes generated herein, were individually transfected into 293T cells (ATCC #CRL-3216). Forty-eight hours post-transfection, cells were frozen, thawed, vortexed, and spun at 3,000xg to remove large debris. Supernatants were expanded by two passages at MOI <0.05 in BV2 cells, and p2 virus supernatant was filtered (0.2 μm), concentrated by ultra-centrifugation through a 30% sucrose cushion, and titered via plaque assay.

Plaque assays were performed in BV2 cells, similar to previously described methods [20]. Briefly, BV2 cells were grown in 6well plates, infected with serial dilutions of each sample (500ul per well, 1h, RT, on a rocking platform), after which the inoculum was removed and cells were overlaid with 1% methylcellulose in complete DMEM. At two to three days post-overlay, cells were fixed and stained with 20% EtOH / 0.1% crystal violet.

### Incucyte live-cell imaging and analysis

Cells were seeded in a 96-well plate, allowed to form a confluent monolayer (~18-24h), and subsequently infected with MNV or UV-inactivated MNV (MOI=0.5). Mock-infected cells, and cells treated with 1µM staurosporine were used for negative and positive controls, respectively. Apoptosis and cell permeabilization were assessed via addition of Incucyte Annexin V Green and Cytox Red reagents (EssenBioscience, Ann Arbor, MI), respectively, following manufacturer protocols. Phase contrast and fluorescent images were collected with a 10x objective using the IncuCyte Dual Dolor ZOOM (EssenBioscience) imaging system at 30minute intervals over a 72h time course, maintained at 37°C and 5% CO_2_. Images were analyzed using Incucyte 2016A software (EssenBioscience) after Top-Hat background subtraction, with a 20µm radius.

### Mice, infections, and tissue collection

*Stat1^−/−^* (*Stat1^tm1Dlv^*) mice were originally obtained from the Jackson Laboratories (stock #012606). *Ifnlr1^−/−^* (generated from *Ifnlr1^tm1a(EUCOMM)Wtsi^*) and *Ifnar1^−/−^* (*Ifnar1^tm1Agt^*) were originally obtained from Washington University in St. Louis. *Ifnlr1^−/−^* and *Ifnar1^−/−^* were bred to generate *Ifnlr1*^−/−^/*Ifnar1*^−/−^ mice (DKO) and *Ifnlr1^+/-^*/*Ifnar1^+/-^* (Dhet). Dhet and DKO crossbreeding yielded littermate mice deficient in neither (Dhet), both (DKO), or a single (*Ifnlr1^+/-^*/*Ifnar1^−/−^* or *Ifnlr1^−/−^*/*Ifnar1^+/-^*) IFN receptor gene(s); these littermates were used for experiments to control for variability between litters.

All mice (7-9weeks old) were perorally infected with 1e6 PFU in a 25μL volume administered by pipet, with the exception of experiments in Figure 5 where 1e6 or 1e7 PFU (as indicated in figure) was administered by oral gavage in a 100μL volume. An equal proportion of males and females were maintained among experimental groups to mitigate results due to any unidentified sex-dependent variables. Same-sex mice, infected with the same MNV strain, were often co-housed for acute infection analyses (less than three days), but singly-housed for extended experiments to prevent continued transmission between animals.

Tissue samples, or a single fecal pellet, were collected in 2ml cryovials with 1mm zirconia/silica beads (Biospec, Bartlesville, OK), frozen in a dry ice/EtOH bath and stored at −80C. For plaque assay or RT-qPCR analyses, either 1ml DMEM or 1ml RiboZol (VWR) was added, respectively, and tissues were homogenized via bead-beating (1min, RT) prior to continued analysis.

### Ethics Statement

All mice were bred on C57BL/6 background and maintained in specific-pathogen-free barrier facilities at Oregon Health & Science University under animal protocols approved by the institutional animal care and use committee at Oregon Health & Science University (protocol # IP00000228) according to standards set forth in the *Animal Welfare Act*.

### RNA, cDNA, and RT-qPCR

RNA from tissues was extracted via RiboZol (VWR) protocols, and RNA from cells was extracted with the ZR Viral RNA kit (Zymoresearch, Irvine, CA). DNA contamination was removed with Turbo DNase (ThermoFisher), and 1ug of RNA was used to generate cDNA with the ImpromII reverse transcriptase (Promega, Madison, WI). Quantitative PCR was performed using PerfeCTa qPCR FastMix II (QuantaBio, Beverly, MA), and the following oligo/probes were used for detection: MNV genomic RNA – Forward primer CACGCCACCGATCTGTTCTG, Reverse primer GCGCTGCGCCATCACTC, and Probe TGCGCTTTGGAACAATGG; RPS29 – Forward primer GCAAATACGGGCTGAACATG, Reverse primer GTCCAACTTAATG AAGCCTATGTC, and Probe CCTTCGCGTACTGCCGGAAGC.

### Cell Lysates and Western blot

BV2 cells were grown and infected in 12well plates (2 e5 cells/well). Cells were lifted with trypsin/0.25% EDTA (Gibco, ThermoFisher), washed with 1X cold PBS, resuspended in 100 μL NP40 lysis buffer (Alfo Aesar, Havervill, MA) with protease inhibitor cocktail (Sigma, St Louis, MO), incubated on ice (15min with intermittent mixing), cleared via centrifugation (1min, 4C, full-speed), and diluted 1:1 with 2x SDS loading buffer (0.125 M Tris [pH 6.8], 4% SDS, 20% glycerol, 0.004% bromophenol blue, 10% beta-mercaptoethanol). Lysates were run on Novex Tris-Glycine gels (ThermoFisher), transferred to 0.2 μm PVDF (BioRad, Hercules, CA), blocked with 5% nonfat dry milk, and probed with the following antibodies (Ab): caspase 3 (CST), PARP (CST), beta-Actin pAb (Invitrogen), MNV NS1/2 (a gift from Dr. Vernon Ward, U. of Otago, Australia), or MNV ProPol (a gift form Dr. Kim Green, NIH). Signal was detected using goat, anti-rabbit IgG-HRP (Jackson ImmunoResearch), developed with ECL substrate (Pierce, ThermoFisher), and imaged with an ImageQuant LAS4000 (GE Healthcare, Little Chalfont, UK).

### Intestinal epithelial cell collection

Epithelial fractions were prepared by non-enzymatic striping as previously described [31]. Mice were euthanized and distal small intestine, cecum, and colon (regions of highest MNV replication) were isolated. Peyer’s patches were removed from small intestine and intestinal tissues were incubated in stripping buffer (10% bovine calf serum, 15 mM HEPES, 5 mM EDTA, 5 mM dithiothreitol [DTT] in PBS) with shaking for 20 min at 37°C. The dissociated cells were collected as the epithelial fraction, consisting predominantly of IECs and directly used for flow cytometry.

### Surface and intracellular staining and flow cytometry

BV2 cells were trypsinized and washed once with 1X PBS, stained with the amine-reactive viability dye, Zombie Aqua (Biolegend, San Diego, CA) (20min, on ice), washed once with cold 2% FBS/PBS, washed once with cold Annexin V binding buffer (CST), and stained with Annexin V-FITC (CST) (15min, RT). Cells were immediately analyzed via flow cytometry on an LSR Fortessa (Becton Dickinson [BD], Franklin Lakes, NJ) with FACSDiva software (BD), and data was analyzed using FlowJo software (FlowJo LLC, Ashland, OR).

IECs isolated as described above were stained with the Zombie Aqua viability dye (Biolegend), Fc receptors were blocked with purified CD16/CD32 pAb (clone93; Biolegend), and cells were probed for surface markers, EpCAM (clone G8.8; Biolegend) and CD45 (clone 30-F11; Biolegend). Cells were subsequently fixed with 2% paraformaldehyde/PBS (10min, RT), and blocked/permeablized in 0.2% Triton X/PBS with 2% normal goat serum (20min, RT). Fixed and permeabilized cells were probed for intracellular viral antigen with 2 distinct antibodies: Rabbit polyclonal anti-NS1/2 (a gift from Dr. Vernon Ward, U. of Otago, Australia) and mouse monoclonal anti-NS1 (clone CM79, a gift from Dr. Ian Clark, U. of Southampton, UK). These unconjugated antibodies were detected with anti-Rabbit Alexa Fluor 647 (Molecular Probes, Eugene, OR), and anti-mouse Dylight 405 (Jackson ImmunoResearch), respectively. Cells were analyzed via flow cytometry, and viable cells that were EpCAM+, CD45-, NS1/2 pAb+, and NS1 mAb+ were labeled as infected epithelial cells.

### RNA *in situ* hybridization via RNAscope®

Ileum was opened longitudinally, pinned apical side up in wax trays, and fixed in 10% neutral-buffered formalin (Sigma) (18-24h, RT), then moved to 70% EtOH. Tissues were embedded in 2% agar (ThermoFisher), the PP was vertically transected, flipped 90 degrees, and re-embedded in 2% agar. Agar blocks were maintained in 70% EtOH prior to being paraffin-embedded. Tissue sections (5um) were cut and maintained at RT with desiccant until processed. RNA *in situ* hybridization was performed using the RNAscope Multiplex Fluorescent v2 kit (Advanced Cell Diagnostics [ACDBio], Newark, CA) per protocol guidelines. Probes were designed by ACDBio for detection of CW3 positive strand RNA (471891-C1), CR6 positive strand RNA (502631-C1), and EpCAM mRNA (418151-C2), and signals were amplified and detected via ACDBio protocols using TSA® plus technology (Perkin Elmer, Waltham, MA). Slides were mounted with ProLong Gold antifade reagent (ThermoFisher), and imaged using a Zeiss ApoTome2 on an Axio Imager, with a Zeiss AxioCam 506 (Zeiss, Jena, Germany).

### Statistical Analyses

Data were analyzed with Prism 7 software (GraphPad Prism Software, La Jolla, CA), with specified tests and significance noted in the figure legends.

## Acknowledgements

The authors would like to thank the OHSU Advanced Light Microscopy Core, Flow Cytometry Core, and Histopathology Shared Resource for technical support, Lena Li for overseeing mouse colonies, and Stephanie Karst for guidance with RNAscope techniques.

## Supporting information

**S1 Video.** M2C-CD300lf cells infected with MNV strain CW3, related to figure 10.

**S2 Video.** M2C-CD300lf cells infected with MNV strain CR6, related to figure 10.

**S3 Video.** M2C-CD300lf cells infected with MNV strain CR6^Δcasp^, related to figure 10.

**S4 Video.** Mock-infected M2C-CD300lf cells, related to figure 10.

**S5 Video.** M2C-CD300lf cells treated with staurosporine, related to figure 10.

